# Scaling of inertial delays in terrestrial mammals

**DOI:** 10.1101/631846

**Authors:** Sayed Naseel Mohamed Thangal, J. Maxwell Donelan

## Abstract

As part of its response to a perturbation, an animal often needs to reposition its body. Inertia acts to oppose motion, delaying the completion of the movement—we refer to this additional elapsed time as inertial delay. As animal size increases, muscle moment arms also increase, but muscles are proportionally weaker, and limb inertia is proportionally larger. Consequently, the scaling of inertial delays is complex. Here, we quantify it using two biomechanical models representing common scenarios in animal locomotion: a distributed mass pendulum approximating swing limb repositioning (swing task), and an inverted pendulum approximating whole body posture recovery (posture task). We parameterized the anatomical, muscular, and inertial properties of these models using literature scaling relationships, then determined inertial delay for each task across a large range of movement magnitudes and the full range of terrestrial mammal sizes. We found that inertial delays scaled with an average of *M*^0.28^ in the swing task and *M*^0.35^ in the posture task across movement magnitudes—larger animals require more absolute time to perform the same movement as small animals. The time available to complete a movement also increases with animal size, but less steeply. Consequently, inertial delays comprise a greater fraction of swing duration and other characteristic movement times in larger animals. We also compared inertial delays to the other component delays within the stimulus-response pathway. As movement magnitude increased, inertial delays exceeded these sensorimotor delays, and this occurred for smaller movements in larger animals. Inertial delays appear to be a challenge for motor control, particularly for bigger movements in larger animals.

## Introduction

Independent of animal size, a fast response time is important to an animal’s survival. A tiny shrew needs to react quickly to escape from a predator, and a massive elephant needs to recover quickly from a loss of balance to prevent a fall. Response time—measured as the total delay between the onset of a perturbation and the completion of the movement of the body—is not just important for relatively rare escapes and falls, but also for more common motor control tasks. This is because even small time delays can destabilize feedback control, requiring animals to have compensatory neuromechanical strategies [1–4]. Response time is relevant to the control of movement both in terms of its absolute duration and its duration relative to the available movement time. For example, the absolute duration of response time matters to avoid a snakebite, which can be equally deadly for small and large animals alike. And the relative response time matters to avoid a trip when galloping, where the response may need to occur within a limb’s swing duration, which takes longer in larger animals [5–7].

Response time is determined, in part, by neuromuscular physiology [2,8–10]. Consider an animal whose foot catches on a vine—the lengthening of the limb muscles activates the stretch reflex, which resists muscle stretch and helps the animal recover its posture [11,12]. This stretch reflex consists of several component delays. There is a sensing delay to detect the stretch and generate action potentials, a nerve conduction delay to conduct the action potentials through the sensory nerve fibres to the spinal cord, and a synaptic delay to process the signal at the sensorimotor synapse. There is another nerve conduction delay as the action potentials are transmitted down the motor nerve fibres, a neuromuscular junction delay to transmit the action potentials across the neuromuscular junction, an electromechanical delay to conduct the action potentials across the muscle and activate the molecular mechanisms involved in cross bridge formation, and a force generation delay for the muscles to develop forces. We group these six component delays together and refer to their sum as “sensorimotor delay” [2].

After the sensorimotor delays, an animal must often physically move its body to a new position. If its muscles could instantaneously generate infinite force, or if the body and its segments were massless, an animal could accomplish this instantly. But of course, muscles have finite strength and bodies have inertia. Consequently, the animal’s inertia impedes the acceleration generated by muscles, further delaying response time. We refer to this as “inertial delay”. Unlike synaptic delay or neuromuscular junction delay which are well-approximated as fixed durations that are independent of the situation, inertial delay is a dynamic process whose duration depends on the movement task and magnitude. It depends on the task because the time to swing a limb to a new position from rest may be different from that required to reject a push to the torso to avoid a fall. It also depends on the magnitude of the required movement because, all else being equal, less time is required to accomplish small adjustments to the body’s position and velocity than large adjustments. Control theorists typically distinguish between effects of this nature and fixed time delays. We nevertheless use the term “delay” here in an effort to keep the concepts approachable to scientists in a broad range of fields. And we are careful to compare the effects of animal size on inertial delay for a given movement magnitude within a given task.

The scaling of inertial delay depends upon how muscle forces, muscle moment arms, and the body’s inertial properties change with body size. When compared to small animals, larger animals have larger muscles and longer moment arms which increase joint torque, but also heavier and longer limbs which increase moment of inertia [13,14]. These properties don’t scale precisely with simplified scaling rules such as geometric or dynamic similarity [15,16]. Consequently, it is not clear whether allometric scaling of muscle forces and muscle moment arms offset size-dependent increases in inertial properties, or vice versa. A similar principle is evident in the scaling of skeletal stress, where the disadvantages predicted for larger animals when assuming simplified scaling rules are reduced or eliminated by compensatory size-related changes in other factors, such as posture and moment arms [17–19].

Here we seek to understand how inertial delays scale with animal size in terrestrial quadrupedal mammals. We begin by deriving scaling relationships for movement durations. We use these relationships to understand inertial delay in context of the time available to move. Next, we focus our study on two different tasks designed to represent scenarios commonly encountered during animal locomotion. The swing task represents an animal repositioning its limb to control foot placement—modeled as a distributed mass pendulum. The posture task represents an animal recovering its posture after a push forward in the sagittal plane—modeled as a point-mass inverted pendulum. We begin this study by deriving analytical expressions for the scaling of inertial delay by linearizing both of these models and parameterizing them using simplified scaling rules. This helps build intuition for the dependence of inertial delay on task, movement magnitude, muscle force, muscle moment arm, and inertial properties. Then to obtain more realistic estimates for the scaling of inertial delays, we parameterize the complete nonlinear models with measured values from literature and simulate them numerically. We estimate response time as the sum of sensorimotor delays and inertial delays. We then compare it to the available movement time, to gauge whether response times reach magnitudes where they could detrimentally affect motor control.

## 2. Scaling of characteristic movement times

We use characteristic movement times to understand how much time an animal has to respond to a perturbation. We compare response time to these movement times to gauge whether the time required to respond may hinder neural control of movement. Here, we analytically quantify the scaling of two characteristic movement times which we have chosen to approximate the time it would take an animal to fall to the ground, and the time an animal’s leg is in swing phase when running.

As response time becomes longer relative to fall time, it becomes more difficult for an animal to stop a fall and regain balance. To analytically derive the scaling of fall time, consider an animal of mass *M*, falling from the height of its leg *L* to the floor. *F*_*g*_ is the force of gravity, where *g* is the acceleration due to gravity. The equations for acceleration *ÿ*(*t*), velocity 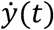 and position *y*(*t*) are:

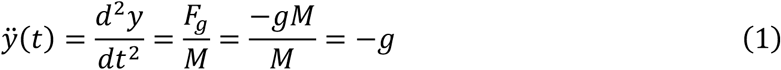

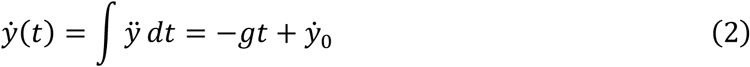

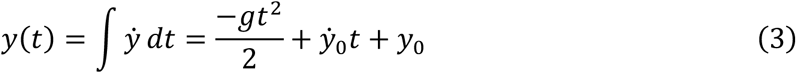

The initial velocity 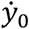 is 0 and initial position *y*_0_ is *L*. Here, we assume that animal morphological features scale with geometric similarity. Two animals are geometrically similar if they have exactly the same shape, even if they are of different sizes. Furthermore, geometric similarity predicts that animal linear features such as leg length scale with *M*^1/3^, surface area features scale with *M*^2/3^, and masses of body segments scale with *M*^1^ if animal density does not change. Then the time *t*_*f*_ required for an animal to fall to the floor is:

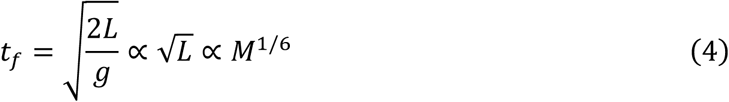

It would take longer for an animal to fall like an inverted pendulum, rather than crumple to the ground as described above, but the dependence on mass would not change. Similar to falling, if response time exceeds the natural time period of the swinging limb, or some fraction of this period, the animal may have difficulty recovering if the swing is perturbed. We used the natural time period of a pendulum with the properties of an animal limb as a proxy for swing duration [20,21]. Assuming geometric similarity, the natural time period *t*_*s*_ of a pendulum scales as:

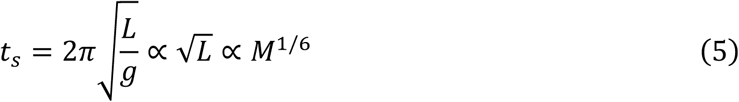

Calculations based on values reported in the literature estimate that swing duration at the trot-gallop transition speed scales with *M*^0.14^ while swing duration at maximum sprint speeds scales with *M*^0.13^ [2]. Thus, characteristic movement times scale approximately with *M* ^1/6^ based on both theoretical considerations and empirical measurements. We use these characteristic movement times to normalize response time and calculate relative delay, a measure of how long response time is when compared to the time available to complete the response.

## 3. A simple model of inertial delay

### 3.1 Model

To obtain theoretical estimates for the scaling of inertial delays, we first consider a simple pendulum operating in the horizontal plane without the effect of gravity (Fig 1). These equations of motion are linear, allowing us to analytically derive the scaling of inertial delays. These estimates will validate subsequent numerical simulations and give us intuition about how various factors contribute to inertial delays. This system is an angular version of the sliding block model and can be analytically described as a double integrator—a simple and well-studied dynamical system [22,23].

**Fig 1.**
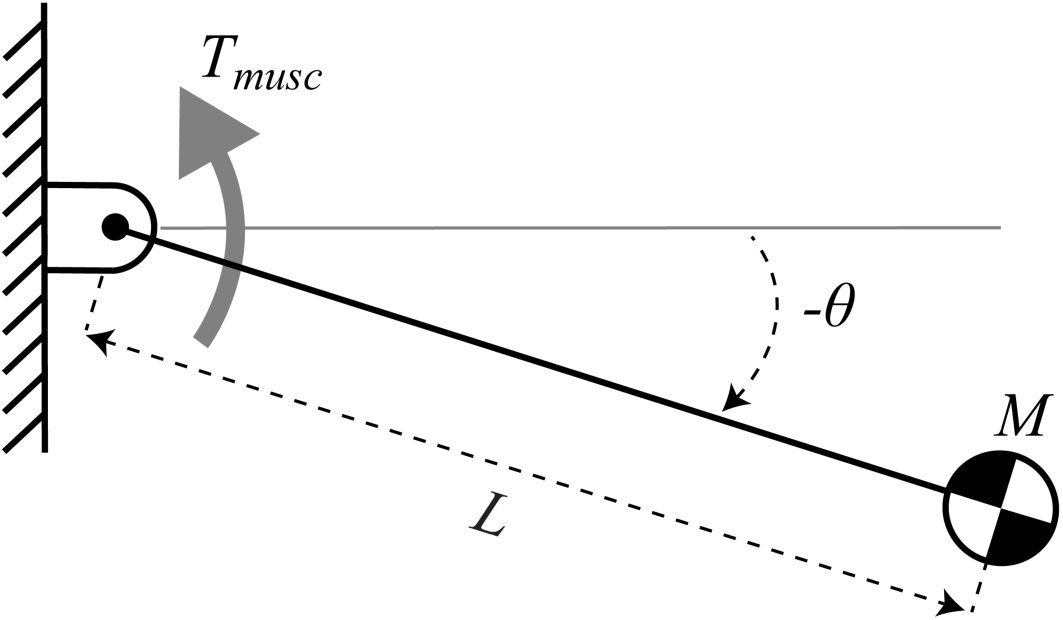
Simple model for inertial delay. A pendulum of rod length *L* and point-mass *M* rotates about a pin joint actuated by torque *T*_*musc*_. *θ* is the angle from the horizontal with positive angles in the counter-clockwise direction. This model ignores gravity and assumes the rod is massless.

### 3.2 Scaling of model parameters

Assuming the pendulum scales with geometric similarity, its length would change with *M*^1/3^, mass with *M*^1^ and moment of inertia (*ML*^2^) with *M*^5/3^. We consider two scenarios for the scaling of maximum muscle force. In the first scenario, we assume that muscle force maintains dynamic similarity between animals of different sizes by scaling force in direct proportion to animal mass: *F*_*musc*_ ∝ *M*^1^ [15]. In the second scenario, we instead assume muscle force scales with cross sectional area, which we refer to as muscle stress similarity: *F*_*musc*_ ∝ *M*^2/3^ [17]. In this model, the applied muscle torque *F*_*musc*_ is the product of a constant muscle moment arm *R*_*musc*_ and the muscle force.

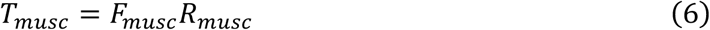

We assume that the muscle moment arm scales with geometric similarity (*M*^1/3^).

### 3.3. Analytical derivation for the swing task

The swing task represents an animal repositioning its swing leg to control foot placement and maintain stability during walking and running [24–28]. For the swing task, the pendulum is required to move from rest at an initial angle *θ*_0_ which we set as the origin, to a final angle *θ*_*f*_ under the control of muscle torque *T*_*musc*_. The fastest way to complete this movement is to apply a constant torque to accelerate from 0 to *θ*_*f*_ /2, then reverse the direction of torque to decelerate and stop at *θ*_*f*_ Since the movement is symmetrical, we consider only the first half from 0 to *θ*_*f*_ /2. The equations for the angular acceleration 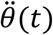, angular velocity 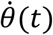 and angle *θ*(*t*) are:

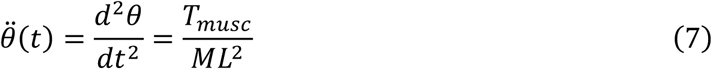

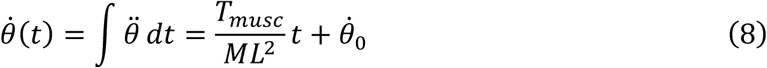

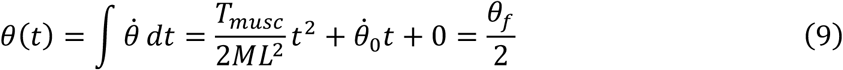

Because the initial velocity 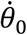 is 0, and our desired final angle is *θ*_*f*_ /2, we can rearrange Eqn 9 to solve for *t*. The total inertial delay is twice this time to account for the time spent in each half of the total movement:

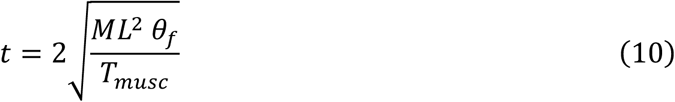

Eqn 10 shows that inertial delay is proportional to the square root of both the movement magnitude (*θ*_*f*_) and the moment of inertia of the pendulum (*ML*^2^), and inversely proportional to the square root of the applied torque. Therefore, doubling muscle torque would only cause an approximately 30% reduction in inertial delay and quadrupling muscle torque would only result in a 50% reduction. These calculations indicate that while inertial delay does depend on actuator limits, increasing muscle torque may not be an effective option to reduce it.

Next, we consider our two different scenarios for the scaling of muscle force. If muscle force scales with dynamic similarity (*F*_*musc*_ ∝ *M*^1^), we can determine the scaling of inertial delay by substituting Eqn 6 into Eqn 10:

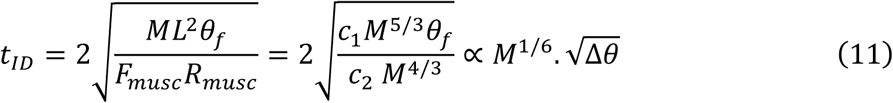

where *c*_1_ and *c*_2_ are constants of proportionality. Since *θ*_0_ is 0, we can substitute *θ*_*f*_ with Δ*θ* = (*θ*_*f*_ − *θ*_0_), representing the movement magnitude.

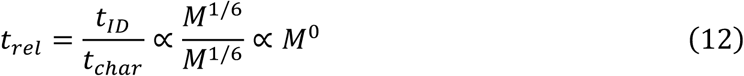

where *t*_*ID*_ is the inertial delay, *t*_*char*_ is the characteristic movement time, and *t*_*rel*_ is the relative delay. Therefore, if muscles produce forces proportional to their mass, inertial delay will scale with the same exponent as characteristic movement times (Eqns 4,5), and relative delay would be independent of animal size. As much as animals are like this simple model with its assumptions, large and small animals would be dynamically similar in their response to disturbances and relative delay would not change with animal size. In this situation, inertial delay would not disproportionately burden larger animals.

Instead, if muscle forces scale with cross sectional area (*F*_*musc*_ ∝ *M*^2/3^), getting relatively weaker with increases in size, inertial delay scales as:

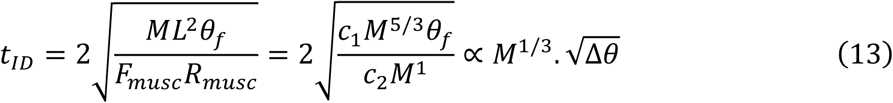

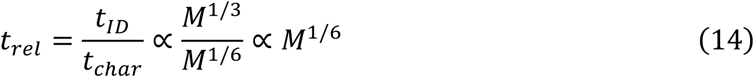

In contrast to dynamic similarity, relative delay will grow with animal size proportional to *M*^1/6^, when muscle force grows only in proportion to cross sectional area, penalizing larger animals. This highlights the importance of muscle force scaling in determining the effect of size on inertial delay.

### 3.4 Analytical derivation for the posture task

The posture task models a standing animal recovering its balance after being perturbed [29–32]. We represent the standing quadruped with a pendulum, which starts from an initial position and has an initial clockwise velocity in the sagittal plane due to a perturbation pushing it forward. We define inertial delay as the time required for muscle torque to return the pendulum to rest back at the initial position after recovering from the perturbation. We again use the simple model (Fig 1) and ignore the effects of gravity. For this task, the movement is not symmetrical. To analytically derive the equations for inertial delay in the posture task, it is convenient to break down the movement into three phases: A, B and C. In phase A, the pendulum starts at an initial position with a clockwise velocity due to the perturbation. We then apply a counter-clockwise torque to decelerate the pendulum and reject the velocity perturbation, stopping at a clockwise angle. In phase B, we continue to apply the counter-clockwise torque, accelerating the pendulum from rest with a counter-clockwise velocity as it moves back towards its initial position. In phase C, we switch the torque direction again so that a clockwise torque now decelerates the pendulum and brings it to rest at the initial position, thereby completing the response to the velocity perturbation. We describe the analytical derivation below.

In phase A, the pendulum starts from an initial angle _*A*_*θ*_0_ with an initial clockwise angular velocity 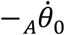. We define _*A*_*θ*_0_ to be the origin with value 0, and counter-clockwise movements to be positive. We then apply a counter-clockwise torque *T*_*musc*_ that brings the pendulum to rest at a final position −_*A*_*θ*_*f*_. The equations for angular acceleration 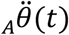, angular velocity 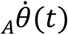 and angle _*A*_*θ*(*t*) are:

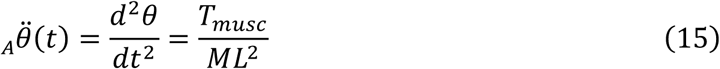

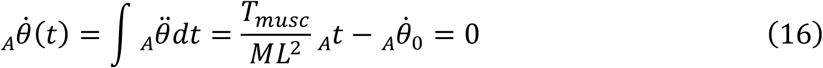

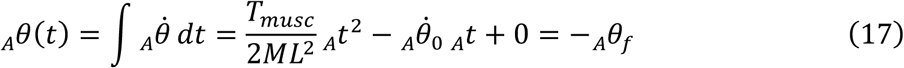

Rearranging Eqn 16, we can express the duration of phase A (_*A*_*t*) in terms of the initial velocity of the perturbation:

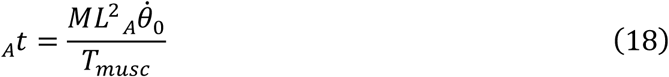

Substituting Eqn 18 into Eqn 17 and simplifying gives us the final angle of phase A (_*A*_*θ*_*f*_) in terms of the initial velocity 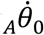:

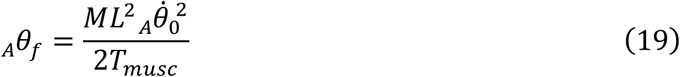

In phases B & C, the pendulum is brought back to rest at the origin from the end position of phase A (−_*A*_*θ_f_*). The movements in phases B and C are equal and opposite, with a counter-clockwise torque initially accelerating the pendulum from rest to a position of −_*A*_*θ_f_* /2 in phase B, followed by a clockwise torque decelerating the pendulum over the same angular distance to rest at the origin in phase C. Therefore, we only need to evaluate the time required for phase B, since the time required for phase C will be the same. The subtask to be accomplished in phases B and C is the same as that of the entire swing task in section 3.3—begin at rest, move through some angular displacement, and end at rest. The additional feature of the posture task is that the initial velocity perturbation determines the subsequent angular displacement. The equations for the angular acceleration 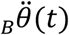, angular velocity 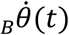 and angle _*B*_*θ*(*t*) are:

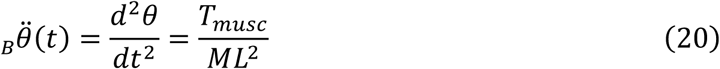

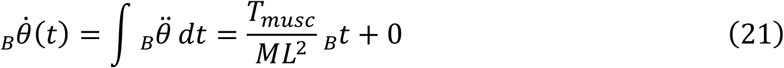

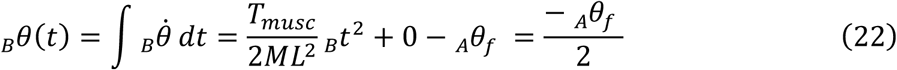

Simplifying Eqn 22 gives:

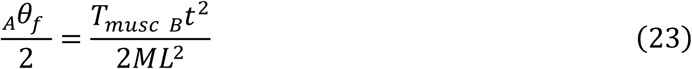

Substituting the final angle of phase A (_*A*_*θ_f_*) from Eqn 19 into Eqn 23 and solving for the duration of phase B (_*B*_*t*) gives:

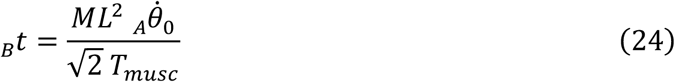

Solving for the total time for the whole motion (phases A, B and C) gives:

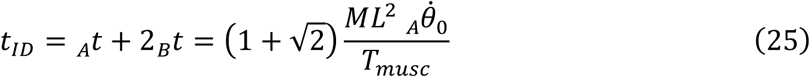

Similarly to Eqn 10 for the swing task, the analytical derivation in Eqn 25 gives us insight into how the various factors contribute to inertial delay (*t*_*ID*_) for the posture task. It predicts that the inertial delay during posture recovery after a perturbation is directly proportional to the perturbation size 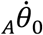 and inversely proportional to the muscle torque *T*_*musc*_.

Since larger animals have heavier bodies, longer limbs and larger muscles, we scaled the size of the perturbation with animal mass to evoke responses with similar relative magnitude. To do this, we express the initial angular velocity of the pendulum, representing the applied perturbation, in terms of linear velocity:

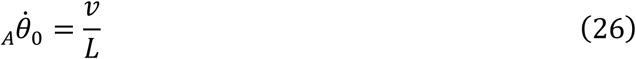

where *v* is the linear velocity caused by the initial perturbation and *L* is the length of the pendulum. We perturbed each model using an initial linear velocity scaled based on a constant dimensionless velocity *v*_*ND*_ [33]:

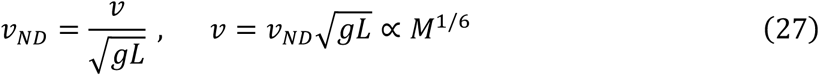

Substituting the values for *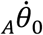* from Eqn 26 and *T*_*musc*_ from Eqn 6 into Eqn 25 and assuming muscle force scales with dynamic similarity (*F*_*musc*_ ∝ *M*^1^) predicts that the total time for the posture task scales as:

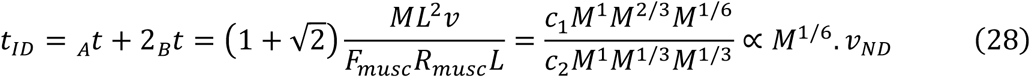

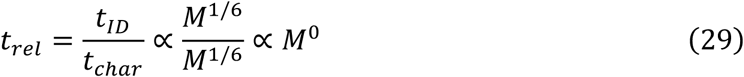

If instead muscle force scales with cross sectional area (*F*_*musc*_ ∝ *M*^2/3^), the total time for the posture task would scale as:

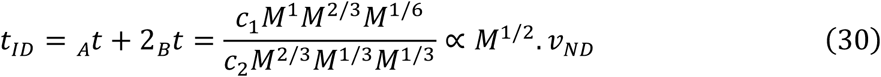

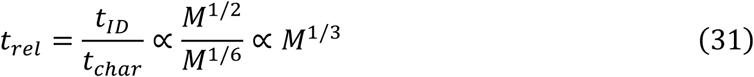

Similar to our conclusions from Eqn 11 in the swing task, we find that if muscles produced forces proportional to their mass (as in dynamic similarity), both inertial delays and characteristic movement times would scale with *M*^1/6^. This would result in constant relative delays regardless of animal size. However, if muscles produced forces proportional to their cross-sectional area, inertial delay would scale with *M*^1/2^ in absolute time and *M*^1/3^ when expressed relative to movement time.

The effect of size on inertial delay depends on the task. The effect of size under muscle stress similarity is steeper in the posture task (Eqn 30) than what we found in the swing task (Eqn 13). An additional difference is the effect of movement magnitude—inertial delay increases in direct proportion to the size of the velocity perturbation in the posture task, and only with the square root of the angular displacement in the swing task. In sections 4 and 5, we use computer simulations of nonlinear biomechanical models, parametrized by actual measurements from literature, to refine our estimates for the scaling of inertial delays.

## 4. Swing task

### 4.1 Model

We modeled the swing task as a distributed mass pendulum actuated by muscle torque (Fig 2). We defined inertial delay for this task as the time required to swing the pendulum from rest at an initial clockwise angle to rest at a final counter-clockwise angle, with identical angles in the clockwise and counter-clockwise direction. Unlike our simple model, we included the effects of gravity, did not assume a point mass, and did not linearize the equation of motion. The motion of the pendulum is described by:

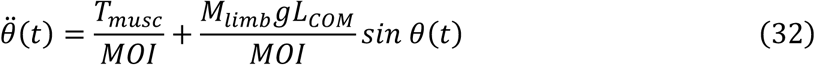

where *T*_*musc*_ is the muscle torque, *L*_*COM*_ is the distance from the pendulum pivot to limb center of mass, *M*_*limb*_ is the mass of the limb, and *MOI* is the moment of inertia of the forelimb about the shoulder joint (Fig 2a). We applied the control torque in a bang-on bang-off profile from +*T*_*musc*_ to −*T*_*musc*_ to determine a lower bound for inertial delay by ignoring realistic muscle actuation dynamics. In this scenario, inertial delay represents the minimum movement time possible, and is limited only by maximal torque (Fig 2b top panel).

**Fig 2.**
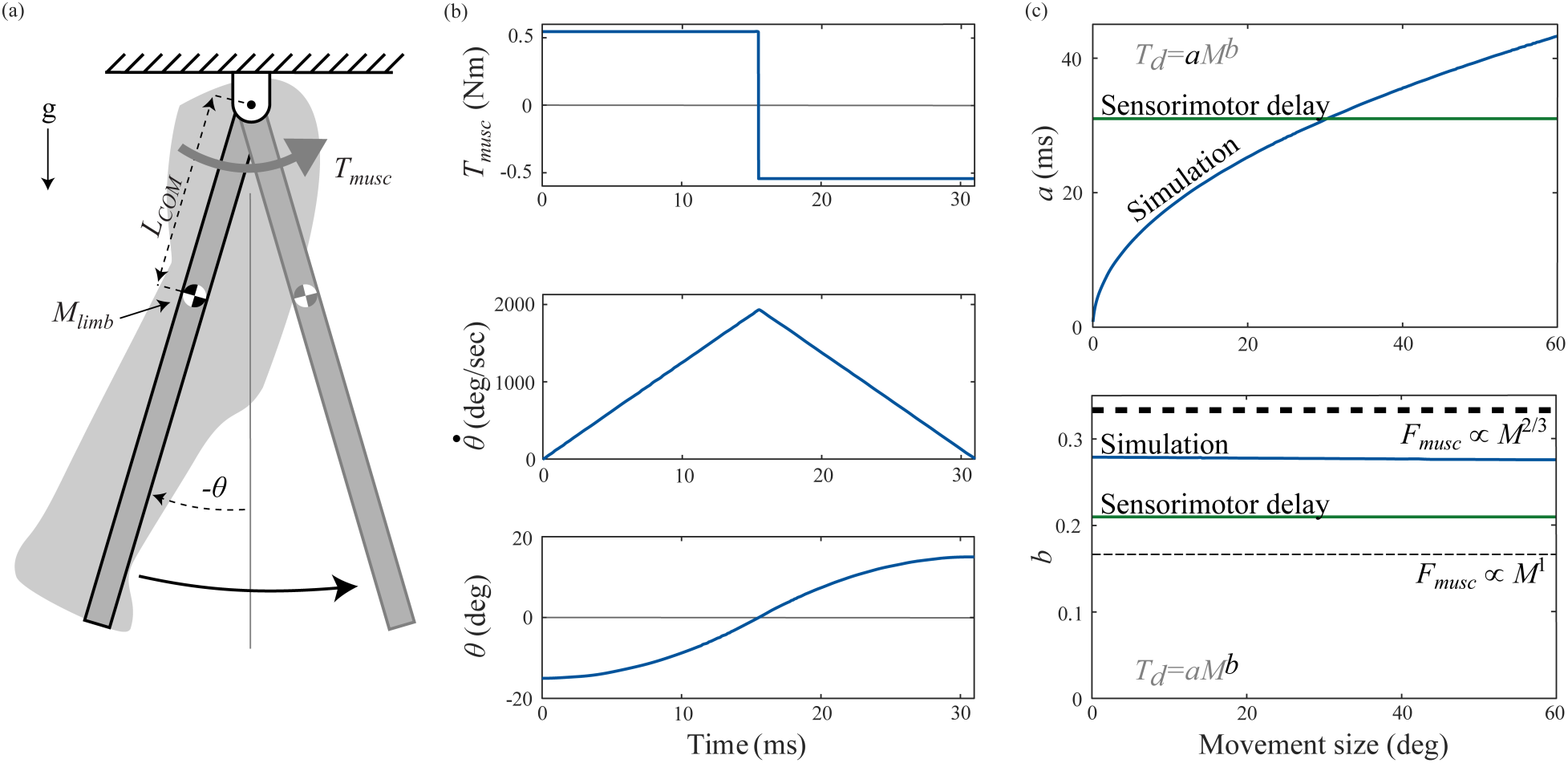
Swing task. The model represents repositioning of the swing leg during locomotion. (a) We modeled the swing task as a distributed mass pendulum actuated by muscle torque *T*_*musc*_. Our model incorporates the distance from the pendulum pivot to limb center of mass (*L*_*COM*_), mass of the limb (*M*_*limb*_), and moment of inertia of the forelimb about the shoulder joint (*MOI*). (b) Angle (*θ*), angular velocity 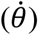, and torque (*T*_*musc*_) profiles in the swing task for a one kg animal for a 30 degree movement, the movement magnitude for which inertial delay equals sensorimotor delays in a one kg animal. (c) Variation in coefficient *a* and exponent *b* of the power law for inertia delay from numerical simulations with movement magnitude (dark blue), sensorimotor delays (dark green), and theoretical predictions for the inertial delay exponent based on scaling of muscle force with cross sectional area ∝ *M*^2/3^ (thick dashed line) and dynamic similarity ∝ *M*^1^ (thin dashed line).

### 4.2 Scaling of model parameters

Table 1 summarizes the scaling relationships we used for our swing task parameters. We used scaling equations for forelimb mass, length, distance from shoulder joint to limb center of mass, and moment of inertia from Kilbourne and Hoffman [14]. We used scaling equations for triceps muscle mass, muscle length, and moment arm from Alexander et al. [13]. We assumed that the triceps is the main muscle flexing the shoulder joint in quadrupeds, because it is a prime mover for this action and because it is the only shoulder muscle for which all the values necessary to compute the scaling of muscle torque are available. Using values for the entire triceps is a further simplification, as only one of the three heads of the triceps move the shoulder [29]. We assumed that parameters for the shoulder extensor muscles scale in the same way as those for the triceps.

**Table 1.** Swing task scaling parameters and their confidence intervals.

| Parameter | Coefficient (*a*) |  |  | Exponent (*b*) |  |  |
| --- | --- | --- | --- | --- | --- | --- |
|  | Value | 95% CI |  | Value | 95% CI |  |
| Forelimb inertial properties [14] |  |  |  |  |  |  |
| Mass (kg) | 5.82×10<sup>-2</sup> | 4.61×10<sup>-2</sup> | 7.34×10<sup>-2</sup> | 1.00 | 0.93 | 1.08 |
| COM length (m) | 5.64×10<sup>-2</sup> | 4.98×10<sup>-2</sup> | 6.38×10<sup>-2</sup> | 0.36 | 0.32 | 0.40 |
| MOI (kg m<sup>2</sup>) | 2.52×10<sup>-4</sup> | 1.61×10<sup>-4</sup> | 3.95×10<sup>-4</sup> | 1.75 | 1.60 | 1.89 |
| Triceps muscle properties [13] |  |  |  |  |  |  |
| Mass (kg) | 6.20×10<sup>-3</sup> | 5.54×10<sup>-3</sup> | 6.94×10<sup>-3</sup> | 1.11 | 1.07 | 1.15 |
| Muscle length (m) | 1.87×10<sup>-2</sup> | 1.72×10<sup>-2</sup> | 2.04×10<sup>-2</sup> | 0.33 | 0.29 | 0.37 |
| Moment arm (m) | 8.70×10<sup>-3</sup> | 8.13×10<sup>-3</sup> | 9.31×10<sup>-3</sup> | 0.41 | 0.38 | 0.44 |

To determine muscle torque for each animal size, we first determined muscle volume by dividing the mass of the muscle by a density of 1060 kg/m^3^ [34]. We found the muscle cross-sectional area by dividing its volume by the muscle length, assuming that muscles have a consistent cross-sectional area. Multiplying the cross-sectional area by the isometric force generation capacity of mammalian muscle, estimated at 20 N/cm^2^, gave muscle force [35,36]. Finally, we calculated muscle torque by multiplying the muscle force and its moment arm (Eqn 6).

### 4.3 Simulation

We performed simulations of the swing task for seven animal masses logarithmically spaced from one gram to ten tons, chosen to span the entire size range of terrestrial mammals [37,38]. For each animal mass, we used the scaling relationships from section 4.2 to determine the size-specific parameters for simulation. At each animal size, we varied the initial clockwise angle from 0.01 to 30 degrees to quantify how movement size affected inertial delay. We numerically simulated the swing task using an explicit Runge-Kutta algorithm implemented with MATLAB’s ode45 solver (MATLAB R2017b, The MathWorks, Inc., Natick, MA, USA). We used the solver’s event detection to determine when the pendulum reached zero angle, and taking advantage of the symmetric nature of the problem, switched the direction of the applied torque from counter-clockwise to clockwise. The simulation continued until the solver’s event detection halted the simulation when the pendulum reached zero angular velocity, which occurred when the pendulum reached the same counter-clockwise angle as it had started in the clockwise direction. Fig 2b shows an example simulation. Elapsed simulation time was the inertial delay for each animal size and each initial angle. For each initial angle, we then logarithmically transformed the inertial delay values for the various animal sizes and used least squares linear regression to extract the coefficient and exponent for the scaling of inertial delay [39].

We used Monte Carlo simulations to determine 95% confidence intervals for our results by propagating the uncertainty in the input scaling values for limb inertial properties and muscle properties through to our estimates for inertial delay [40,41]. First, we generated probability distributions for each of the limb inertial and muscle properties in Eqn 32. For the inertial properties, we fit a linear regression model in MATLAB to the log-transformed raw data from Kilbourne and Hoffman, to generate parameters describing the distribution of the coefficient *a* and exponent *b* [14]. For the muscle properties, we did not have access to the raw data so we used the mean and 95% confidence intervals of the scaling parameters to generate t-distributions [13]. We then randomly sampled a single value for each limb inertial and muscle property from their respective probability distribution and used them to simulate our model, generating one scaling coefficient and exponent for inertial delay. We ran 10,000 simulations in this way, obtaining a distribution of coefficients and exponents. Our final 95% confidence intervals are 1.96 times the standard deviations of these distributions.

### 4.4 Results

Our numerical simulations determined that inertial delay scales with an average of *M*^0.28^ for the swing task, across movement magnitudes (Fig 2c). This scaling exponent falls between our two analytical predictions, which assume that muscle force scales either with dynamic similarity *M*^1/6^ (Eqn 11) or with muscle cross sectional area *M*^1/3^ (Eqn 13). The coefficient of inertial delay in our numerical simulations increased with the square root of movement size (Fig 2c top), as predicted by our analytical analysis (Eqn 10). As movement size increased from 1 degree to 60 degrees, the coefficient increased from 5.8 ms (4.0–7.5 ms) to 43 ms (30–57 ms), while the exponent remained fairly steady about 0.28 (0.22–0.34). Here and elsewhere, we report our results as “mean (lower 95 % confidence interval – upper 95% confidence interval)”.

We tested the sensitivity of our numerical results to the applied muscle torque. Varying the torque from half to four times its original value only increased the scaling exponent of inertial delay from *M*^0.276^ to *M*^0.279^. This indicates that our results for the scaling of inertial delay are robust to possible inaccuracies in our estimates for the torque produced by muscles that flex and extend the shoulder joint.

## 5. Posture task

### 5.1 Model

We modeled the posture task as an inverted pendulum that has been pushed forward resulting in an initial body velocity (Fig 3a). The task goal is to apply the correct muscle forces to reject the perturbation and return the inverted pendulum to rest in an upright posture. We defined inertial delay for this task as the time required to move from a vertical position with an initial velocity perturbation in the clockwise direction to rest at the vertical position, under the control of muscle torque. Unlike our simple model of this task, we included the effects of gravity and did not linearize the equation of motion. The motion of this inverted pendulum model is described by:

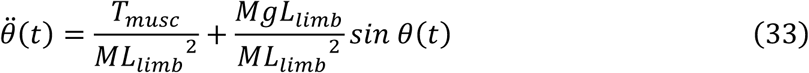

where *T*_*musc*_ is the muscle torque, *L*_*limb*_ is the average length of the forelimb and hindlimb, and *M* is the total mass of the animal. As in the swing task, we applied the control torque in a bang on bang off profile. In this scenario, inertial delay represents the minimum movement time possible, and is limited only by maximal torque.

**Fig 3.**
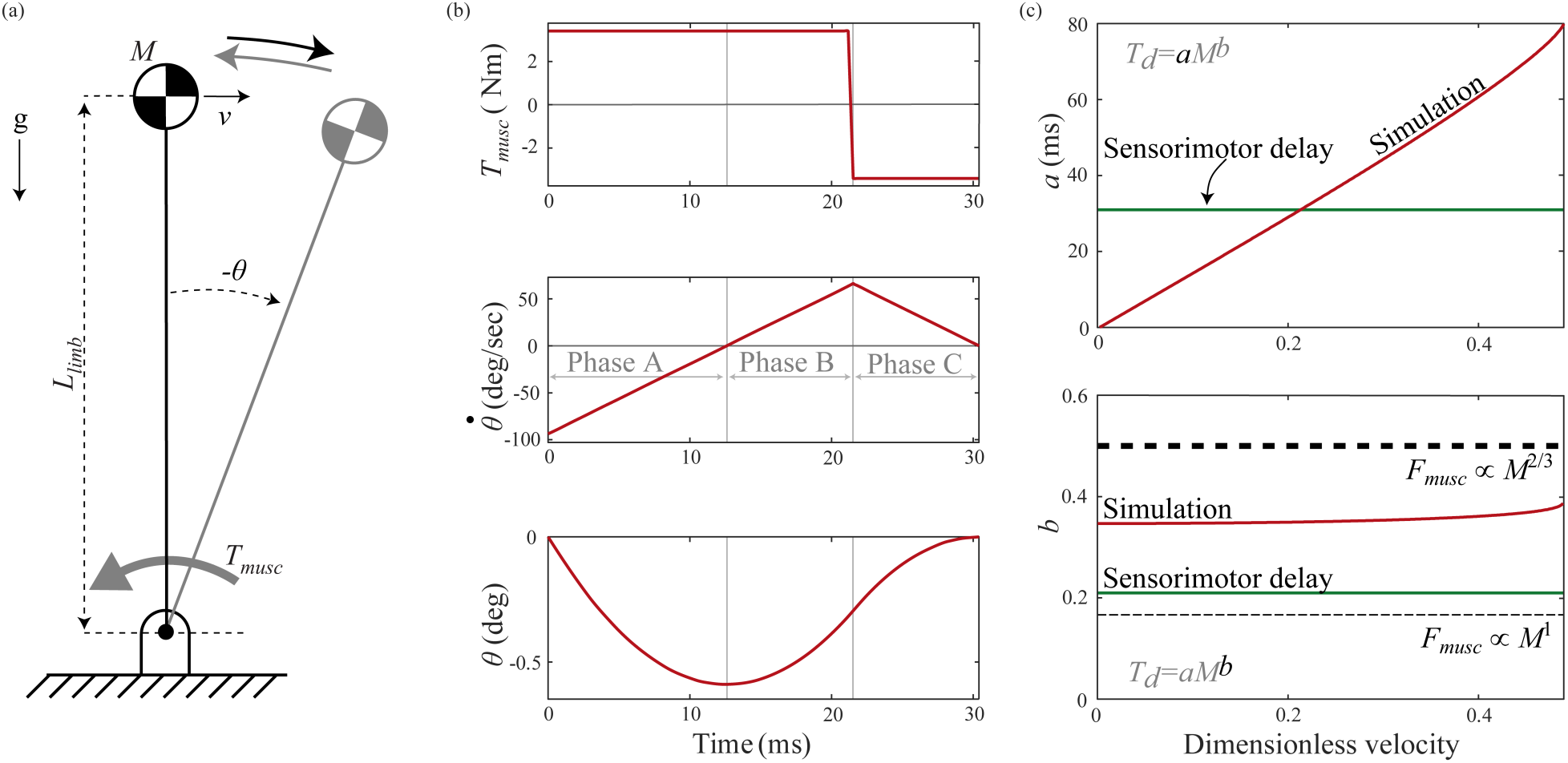
Posture task. The model represents an animal recovering its posture after a perturbation (a) We use an inverted pendulum with a point-mass body and massless rigid legs, pivoting about a ground-mounted pin joint. Our model considers limb length (*L*_*limb*_), mass of animal (*M*), and actuating muscle torque (*T*_*musc*_). (b) Angle (*θ*), angular velocity 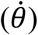, and torque (*T*_*musc*_) profiles in the posture task for a one kg animal for movement of 0.21 dimensionless velocity, the perturbation size for which inertial delay equals sensorimotor delay in a one kg animal. (c) Variation in coefficient *a* and exponent *b* of the power law for inertial delay determined from numerical simulations with perturbation size (red), sensorimotor delays (dark green), and theoretical predictions for the inertial delay exponent based on scaling of muscle force with cross sectional area ∝ *M*^2/3^ (thick dashed line) and dynamic similarity ∝ *M*^1^ (thin dashed line).

### 5.2 Scaling of model parameters

Table 2 summarizes the scaling relationships we used for posture task parameters. We set the length of the inverted pendulum as the average length of the hindlimb and forelimb from Kilbourne and Hoffman, because we wanted the pendulum mass to represent the whole-body center of mass of the animal [14]. In contrast, we set the swing task pendulum length to the length of the forelimb. If we had used the length of the forelimb for the posture task inverted pendulum, our values would increase by 8% or less. We used scaling equations for ankle extensor muscle mass, muscle length, and moment arm from Alexander et al. and computed muscle torque using the steps described in section 4.2 [13]. We assumed that the posture of the animal is controlled by the ankle extensor muscle groups on the four legs by setting *T*_*musc*_ to be four times the torque applied by the ankle extensor group.

**Table 2.** Posture task scaling parameters and their confidence intervals.

| Parameter | Coefficient (*a*) |  |  | Exponent (*b*) |  |  |
| --- | --- | --- | --- | --- | --- | --- |
|  | Value | 95% CI |  | Value | 95% CI |  |
| Limb lengths [14] |  |  |  |  |  |  |
| Forelimb length (m) | 1.61×10<sup>-1</sup> | 1.42×10<sup>-1</sup> | 1.82×10<sup>-1</sup> | 0.38 | 0.34 | 0.42 |
| Hindlimb length (m) | 1.63×10<sup>-1</sup> | 1.47×10<sup>-1</sup> | 1.80×10<sup>-1</sup> | 0.36 | 0.32 | 0.39 |
| Ankle extensor muscle properties [13] |  |  |  |  |  |  |
| Mass (kg) | 5.10×10<sup>-3</sup> | 4.40×10<sup>-3</sup> | 5.92×10<sup>-3</sup> | 0.97 | 0.92 | 1.02 |
| Muscle length (m) | 1.06×10<sup>-2</sup> | 8.98×10<sup>-3</sup> | 1.25×10<sup>-2</sup> | 0.14 | 0.06 | 0.22 |
| Moment arm (m) | 9.40×10<sup>-3</sup> | 8.79×10<sup>-3</sup> | 1.01×10<sup>-2</sup> | 0.38 | 0.35 | 0.41 |

### 5.3 Simulation

We performed simulations of the posture task over the same size range as the swing task. For each animal mass, we used the scaling relationships from section 5.2 to determine the size-specific parameters. At each animal size, we scaled the perturbation size based on dimensionless velocity (Eqn 27) to evoke a proportional response from each animal size. The inverted pendulum can reject the perturbation and return to rest at the vertical position only up to a certain limit—if the initial clockwise velocity is too large, the counter-clockwise torque cannot prevent the inverted pendulum from falling to the ground. The largest perturbation that a 10,000 kg animal could reject and return to vertical was 0.49 dimensionless velocity, so we varied the initial perturbation from 0.01 to 0.49 dimensionless velocity. As with the swing task, we numerically simulated the motion in MATLAB. We used optimization to determine when to switch between the maximum counter-clockwise and clockwise torque magnitudes such that the pendulum reached the original upright posture at the same instant the velocity went to zero. For each perturbation magnitude and animal size, we seeded the optimization with an initial guess of the optimal timing and then used the Trust-region dogleg optimization algorithm (implemented using MATLAB’s fsolve function) [42]. It used repeated model simulations to search for the optimal time to switch torque direction. Fig 3 illustrates a representative optimal solution. Elapsed simulation time was the inertial delay for each animal size and each initial angle. Similar to the swing task, we repeated the simulations and optimizations for a range of animal masses and used least squares linear regression to extract the coefficient and exponent for the scaling of inertial delay. We then used Monte Carlo simulations to estimate the 95% confidence intervals.

### 5.4 Results

Our numerical simulations determined that inertial delay scaled with an average of *M*^0.35^ for the posture task, across perturbation magnitudes (Fig 3c). The exponent again fell between those of the analytical predictions assuming muscle force scaling based on dynamic similarity (*M*^1/6^; Eqn 28) and on muscle cross sectional area (*M*^1/2^; Eqn 30). Varying the perturbation size from 0.01 to 0.49 dimensionless velocity caused the coefficient to increase nearly linearly from 1.5 ms (1.1–1.9 ms) to 81 ms (53–102 ms), while the exponent again remained fairly steady about 0.35 (0.24– 0.46). This linear dependence on perturbation size was captured by our simple model of this task (Eqn 25).

## 6. Inertial delays, sensorimotor delays and response time

In both the swing and posture task, inertial delay increases more steeply with animal size than sensorimotor delay. Previous research in our lab has studied sensorimotor delay in terrestrial mammals of varying sizes and found that it scales with *M*^0.21^ [2]. Here we found that swing and posture task inertial delays scaled with an average of *M*^0.28^ and *M*^0.35^ across perturbation magnitudes, respectively. This does not necessitate that inertial delays always exceed sensorimotor delay because inertial delays also depend on the movement magnitude—for very small position changes and velocity perturbations, inertial delays are far shorter than sensorimotor delay at all animal sizes. But as movement magnitudes increase, there reaches a magnitude at which inertial delay first matches, and then exceeds sensorimotor delay. This occurs at smaller movement magnitudes in larger animals (Fig 4). For the swing task, inertial delay exceeded sensorimotor delay when the limb swung through angles greater than 30*M*^-0.14^ degrees, corresponding to 63 degrees in a five gram shrew and only 9 degrees in a five ton elephant. Shrews experience these limb angles only while galloping, but elephants experience them at slower speeds [16]. For the posture task, inertial delay exceeded sensorimotor delay for velocity perturbations greater than 0.21*M*^-0.14^ dimensionless velocity, corresponding to 0.44 in a five gram shrew and only 0.06 in a five ton elephant. For a shrew, this perturbation magnitude is equivalent to its walk-trot transition speed, but for an elephant it is equivalent to a much slower speed [15]. Because day-to-day activities generally involve smaller movements and less extreme perturbations, in most situations sensorimotor delay likely dominates response time for smaller animals while inertial delays dominate for larger animals.

**Fig 4.**
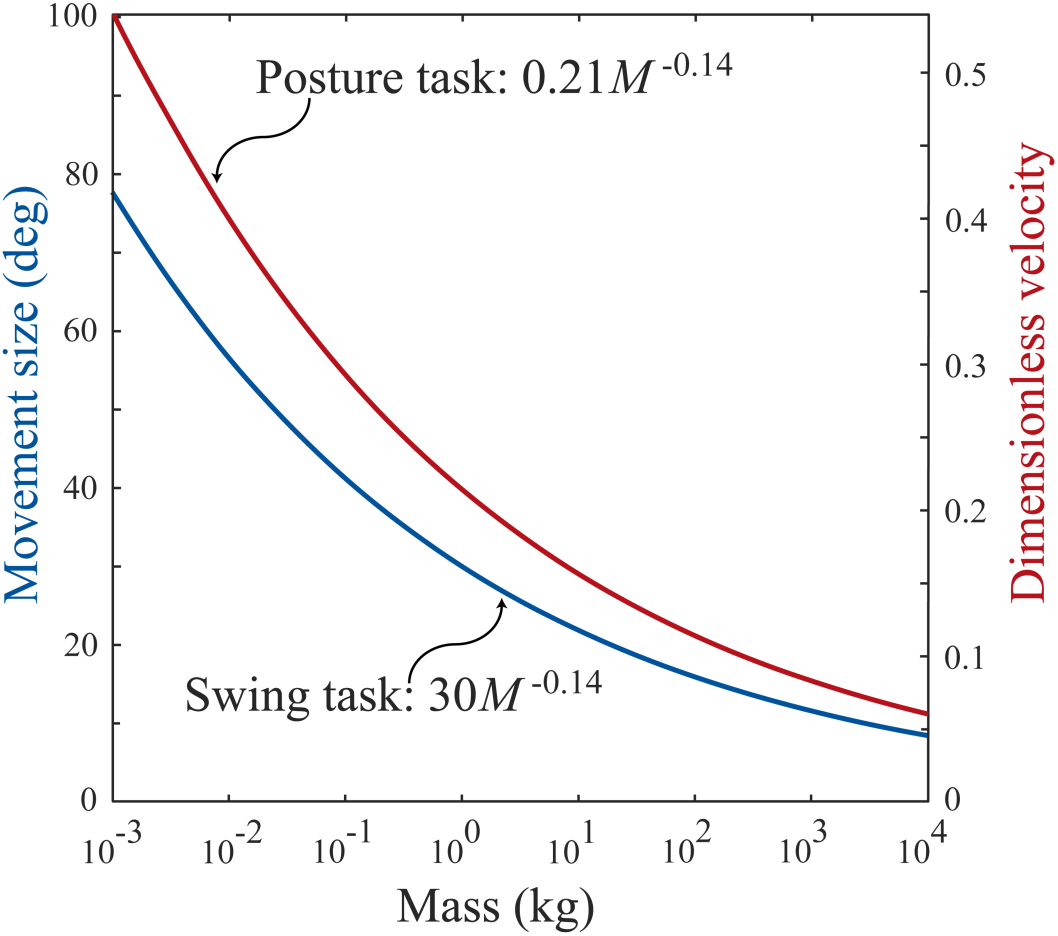
Scaling of movements for which inertial delay equaled sensorimotor delay.

The dependence of both sensorimotor and inertial delays on animal size results in relatively long response times in larger animals. We estimated response time as the sum of sensorimotor delay and inertial delay. If this response time equals or exceeds movement duration, an animal cannot complete the task within the available time. Here, we use swing duration at maximum sprint speed, which scales as 148*M*^0.13^ ms, as a characteristic movement time for fast locomotion [2]. We calculate relative response time as the response time normalized by the available time—it scales as 0.42*M*^0.08^. Inertial delay depends on movement magnitude, and here we assumed a 30 degree swing because at this magnitude, inertial delay matches sensorimotor delay in a 1 kg animal. The fraction of swing duration taken up by sensorimotor delay doubles over seven orders of magnitude of animal mass, while that of inertial delay increases almost six-fold (Fig 5). At maximum running speed, response time requires only about 30% of swing duration for a five gram shrew but about 80% for a five ton elephant. These relatively slower response times in larger animals may hinder their effective control of movement.

**Fig 5.**
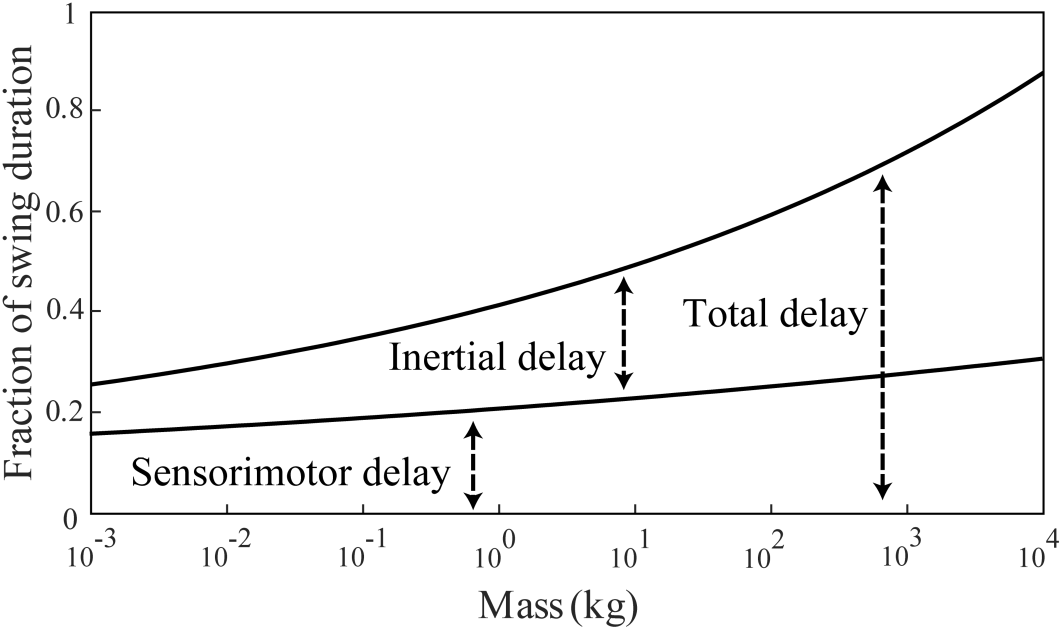
Relative response time. Inertial delay and sensorimotor delay expressed as fractions of swing duration at maximum sprint speed. Inertial delay is shown for a movement of 30 degrees in swing task. At this movement size, inertial delay matches sensorimotor delay in a 1 kg animal.

## 7. Discussion

Here we studied how inertial delays scale with animal size in terrestrial quadrupedal mammals. We defined inertial delay as the component of response time associated with overcoming inertia to move body segments or reject a perturbation and quantified it by modeling two scenarios commonly encountered during animal locomotion. The scaling of inertial delays depended on both the movement task and its magnitude. Over the perturbation magnitudes that we considered, inertial delays scaled with an average of *M*^0.28^ for the swing task and *M*^0.35^ for the posture task, which are both steeper than sensorimotor delays at *M*^0.21^ [2]. We used analytical derivations to show theoretically that if animal muscles could produce forces proportional to an animal’s mass, as required for dynamic similarity, inertial delays would scale at the same rate as characteristic movement times and relative delay would be independent of animal size. However, if muscles only produce forces proportional to their cross-sectional area, relative delay would increase with animal size and disproportionately burden larger animals. Our numerical predictions for the scaling exponent fell between these theoretical predictions indicating that muscle forces that scale more steeply than assumed by stress similarity, and moment arms that scale more steeply than assumed by geometric similarity, partly, but not completely, overcome the increases in inertia with animal size.

Previous work has suggested that animals may be more acutely challenged by long sensorimotor delays than by inertial delays [14]. Our comparison of these two contributors to response time indicates that this is certainly true in all animals when the movement magnitude is small. But for larger movement magnitudes, including magnitudes encountered during day-to-day movements, inertial delay is greater than sensorimotor delay in larger animals (Fig 4). But sensorimotor delays appear to always be important—response time is never entirely dominated by inertial delay (Fig 5). Whether sensorimotor or inertial delays are more challenging to motor control depends on both the movement magnitude and the animal size.

Our study had several important limitations. First, the lack of literature on scaling of muscle properties constrained the accuracy of our estimates for scaling of muscle torque. To our knowledge, only one study reports the scaling of muscle features necessary for determining torques acting about the shoulder and ankle joints in quadrupedal mammals [13]. Second, due to the lack of data for other muscles, we assumed that the triceps and the ankle extensors are the dominant muscles involved in moving their respective joints and that their antagonistic muscles scale similarly. Thirdly, we assumed that the isometric stress produced by mammalian muscle is constant at 20 N/cm^2^ [35], although actual isometric stress values for mammalian muscle vary from 7 to 148 N/cm^2^ [43–45]. We tested the sensitivity of our results to muscle torque for the swing task and found very little effect. This is due to the dependence of inertial delay on the inverse square root of muscle torque, which causes larger torques to give diminishing returns in reduction of inertia delay (section 3.3; Eqn 10). Finally, our models are greatly simplified versions of the rather complex multi-jointed, multi-muscled animal. A more complete model of different size animals, like might be possible with Open-SIMM or a similar approach may provide more realistic estimates of inertial delay [46]. However, we don’t expect that more complete musculoskeletal models would greatly change the identified scaling exponents which were robust to the major simplifications of the analytical models of Section 3 when compared to our nonlinear simulations in Sections 4 and 5.

Our estimate of response time as the sum of sensorimotor delay and inertial delay makes several simplifications. Firstly, we assumed that muscles can produce their maximal forces instantaneously. However, actual muscles have properties that limit their rate of force production, such as activation-deactivation dynamics and force-velocity properties [36,47–50]. Secondly, we assumed that electromechanical delay, force generation delay and inertial delay are distinct. However, these component delays are dynamic processes that overlap [2,49]. Finally, physiological control rarely works in a purely feedforward fashion without sensory feedback. Feedback control is more resilient to unexpected perturbations and to the inherent noise and delays in biological control systems [51,52]. While superior in these regards, it would only slow the response time that we have estimated here—the optimal feedforward control profile operating at the limits to muscle torque yields a response time that is a lower bound on what is possible with feedback control. We suspect that these limitations make our present estimates of response time conservative, and that a refined model or an experimental approach will find response times that exceed movement times, particularly in large animals at fast speeds.

Inertial delays increased significantly with animal size and varied with both movement type and movement magnitude, dominating response time for progressively smaller movements in progressively larger animals. Overall, larger animals face significant delays, both in absolute terms and when considered as a fraction of movement duration. These delays may especially challenge larger animals in situations where they need to react quickly.

## Acknowledgements

We thank David Remy and James Wakeling for insightful comments and suggestions on the project, and the SFU Locomotion Lab for constructive feedback on the manuscript drafts. We are grateful to Heather More for her contribution to figure design, organizational suggestions and manuscript editing.

